# Effect of digital noise-reduction processing on subcortical speech encoding and relationship to behavioral outcomes

**DOI:** 10.1101/2024.10.28.620630

**Authors:** Subong Kim, Mary Schroeder, Hari M. Bharadwaj

## Abstract

Perceptual benefits from digital noise reduction (NR) vary among individuals with different noise tolerance and sensitivity to distortions introduced in NR-processed speech; however, the physiological bases of the variance are understudied. Here, we developed objective measures of speech encoding in the ascending pathway as candidate measures of individual noise tolerance and sensitivity to NR-processed speech using the brainstem responses to speech syllable /da/. The speech-evoked brainstem response was found to be sensitive to the addition of noise and NR processing. The NR effects on the consonant and vowel portion of the responses were robustly quantified using response-to-response correlation metrics and spectral amplitude ratios, respectively. Further, the f0 amplitude ratios between conditions correlated with behavioral accuracy with NR. These findings suggest that investigating the NR effects on bottom-up speech encoding using brainstem measures is feasible and that individual subcortical encoding of NR-processed speech may relate to individual behavioral outcomes with NR.

## 1. Introduction

Modern hearing aids utilize noise-reduction (NR) algorithms to attenuate the noise level to help people with hearing impairment in challenging listening environments [e.g., Bentler and Chiou (2006); Bentler et al. (2008)]. Unfortunately, the NR processing typically used in hearing aids often introduces speech distortion whenever there is spectral overlap between target speech and the background noise [e.g., Arehart et al. (2013); Kates (2008)]. This drawback has been a significant concern for individuals who might perceive speech distortions more sensitively and prefer not to use NR, whereas other users find the benefit of attenuated noise outweighs such distortions and prefer to use NR (Brons, Dreschler, et al., 2014; Brons, Houben, et al., 2014; Neher, 2014; Neher et al., 2014; Neher & Wagener, 2016; Neher et al., 2016). However, little is known about the physiological mechanisms underlying this individual variability. The current study focused on *subcortical* speech encoding in the presence of noise and investigated the effect of NR in the ascending auditory pathway.

Subcortical auditory processing varies even among individuals with normal hearing (Bharadwaj et al., 2022; Bharadwaj et al., 2019; Bharadwaj et al., 2015; Moore, 2008; Picton, 2013; Plack et al., 2014; Ruggles et al., 2011). As a measure of subcortical auditory function, the scalp-recorded auditory brainstem response (ABR) provides non-invasive insights into the ascending auditory system’s ability to encode and process sounds (Felix et al., 2018; Hall, 1992; Krishnan, 2023). Several studies have shown that speech-evoked ABR using brief syllables such as /da/, or ABR to complex sounds (cABR), can be a sensitive tool to probe the fidelity with which an individual’s ascending auditory system encodes the spectrotemporal features of speech sounds (BinKhamis, Léger, et al., 2019; Easwar et al., 2020; Kraus & Nicol, 2005; Nuttall et al., 2015; Skoe & Kraus, 2010), with strong correlations observed between characteristics of speech-evoked ABRs and behavioral outcomes in noise (Anderson & Kraus, 2010; Anderson et al., 2013; Bidelman & Momtaz, 2021; Hornickel et al., 2009; Parbery-Clark et al., 2009). Further, speech-evoked ABR has significant potential as an objective aided outcome measure (Anderson & Kraus, 2013; Easwar et al., 2023; Easwar et al., 2015; Jenkins et al., 2018; Karawani et al., 2018).

Few studies have utilized speech-evoked ABR to evaluate an individual’s physiological reaction to signal processing schemes employed in hearing aids, such as NR. The literature has documented that the effects of background noise on the subcortical processing of speech sounds significantly differ among individuals [e.g., Parbery-Clark et al. (2009); Song et al. (2011); Wong et al. (2007)]. Nevertheless, in investigating the physiological reaction to NR-processed speech, an additional layer of complexity arises because of potential spectral distortions induced by NR, in addition to attenuated noise levels (Kim et al., 2024; Kim, Schwalje, et al., 2021; Kim et al., 2022). Recent studies investigated these mixed effects of NR on *cortical* representations of target speech in the presence of noise (Alickovic et al., 2020; Alickovic et al., 2021; Kim et al., 2022). However, ABR measures have the advantage of evaluating the physiological processing of speech sounds in the early sensory portion of the auditory pathway while being relatively unaffected by top-down processes such as attentional modulation (Figarola et al., 2023; Varghese et al., 2015), and thus more readily applicable in the audiology clinic (BinKhamis, Léger, et al., 2019; Jafari et al., 2015; Rocha-Muniz et al., 2014).

The current study utilized brainstem responses to speech syllable /da/ to investigate the effect of NR on speech encoding in the ascending auditory pathway and relationship to behavioral outcomes. We hypothesized that temporal and spectral characteristics of speech-evoked ABR are sensitive to the addition of noise and NR processing and that such subcortical index relates to behavioral outcomes with NR.

## 2. Results

### 2.1 Response-to-response Correlations

Correlation coefficients calculated from the consonant portions of brainstem responses (5-60 ms) were compared (**Figure 1A**) across different stimulus conditions. Paired *t*-tests showed that a greater degree of similarity was revealed between brainstem responses in the NR condition compared with quiet than the responses in the noise condition compared with quiet (*t*(25) = -3.93, *p* < 0.001; noise-to-quiet: mean = 0.14, SD = 0.083; NR-to-quiet: mean = 0.25, SD = 0.096) (**Figure 1B**). This result suggests that the effect of NR on the brainstem response can be robustly quantified using response-to-response correlation metrics and that NR reduces the degradative effect of noise on the responses.

**Figure 1.**
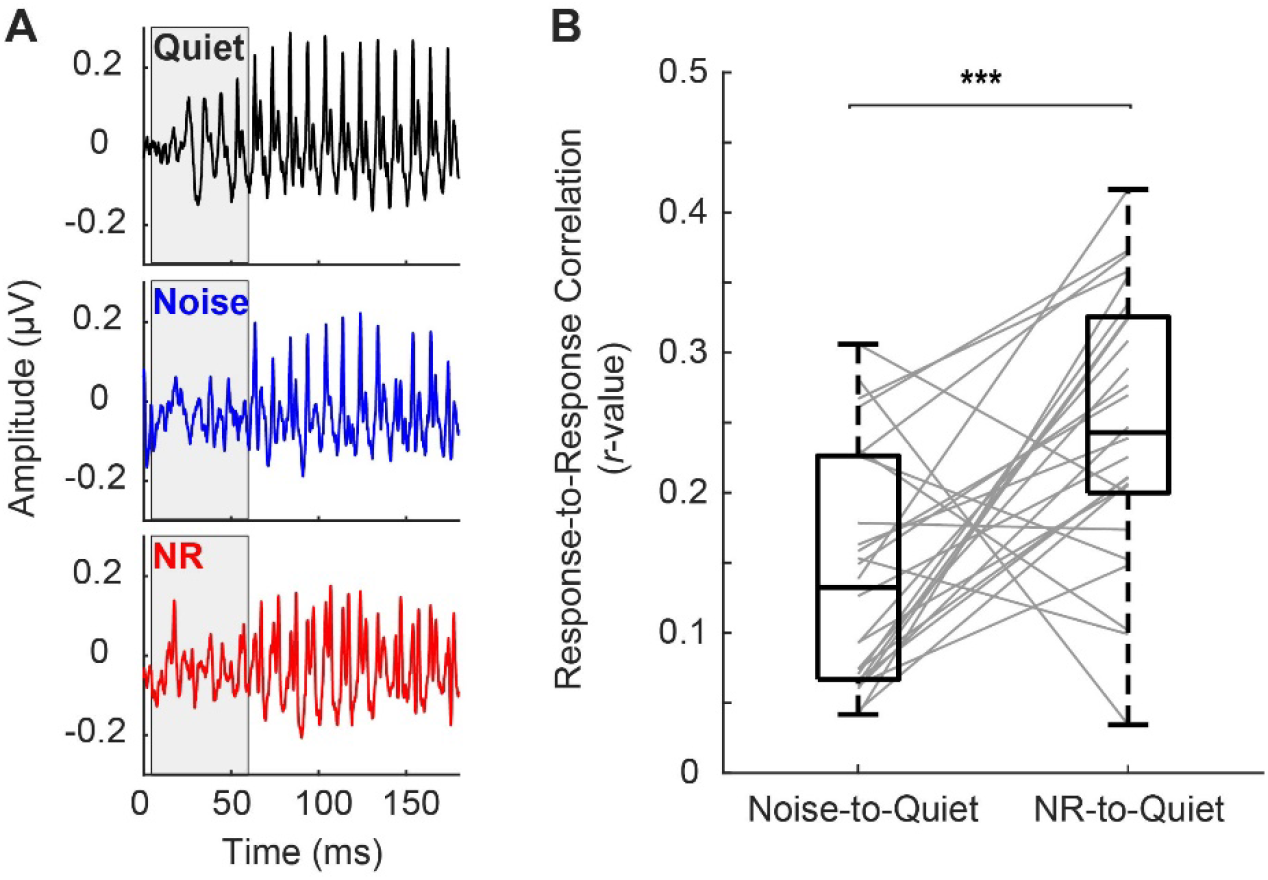
**A.** Auditory brainstem response waveforms in three conditions with shaded regions indicating the consonant portion of the response (5-60 ms). **B**. The noise reduction (NR)-to-quiet correlation was significantly greater than the noise-to-quiet correlation (t(25) = -3.93, p < 0.001), suggesting the role of NR in limiting the degradative effect of noise. ^***^: significant at p < 0.001.

### 2.2 Spectral Encoding

The amplitude spectrums resulting from the fast Fourier transform performed on the vowel portions of the response (60-180 ms) were compared (**Figure 2A**) across different conditions. For f0, paired *t*-test results showed that spectral amplitude in the NR condition compared with quiet was significantly greater than the amplitude in the noise condition compared with quiet (*t*(25) = -3.57, *p* = 0.0015; noise-to-quiet ratio: mean = 0.78, SD = 0.35; NR-to-quiet ratio: mean = 1.11, SD = 0.54) (**Figure 2B** left panel). For upper harmonics (H2-H10), no significant difference was revealed between the NR-to-quiet and noise-to-quiet amplitude ratios (*t*(25) = -1.84, *p* = 0.078; noise-to-quiet ratio: mean = 0.77, SD = 0.095; NR-to-quiet ratio: mean = 0.82, SD = 0.12) (**Figure 2B** right panel). This result suggests that spectral amplitude ratios of brainstem responses between conditions are sensitive to NR effects (especially f0 amplitude) and that NR processing can lead to enhanced spectral encoding in noise.

**Figure 2.**
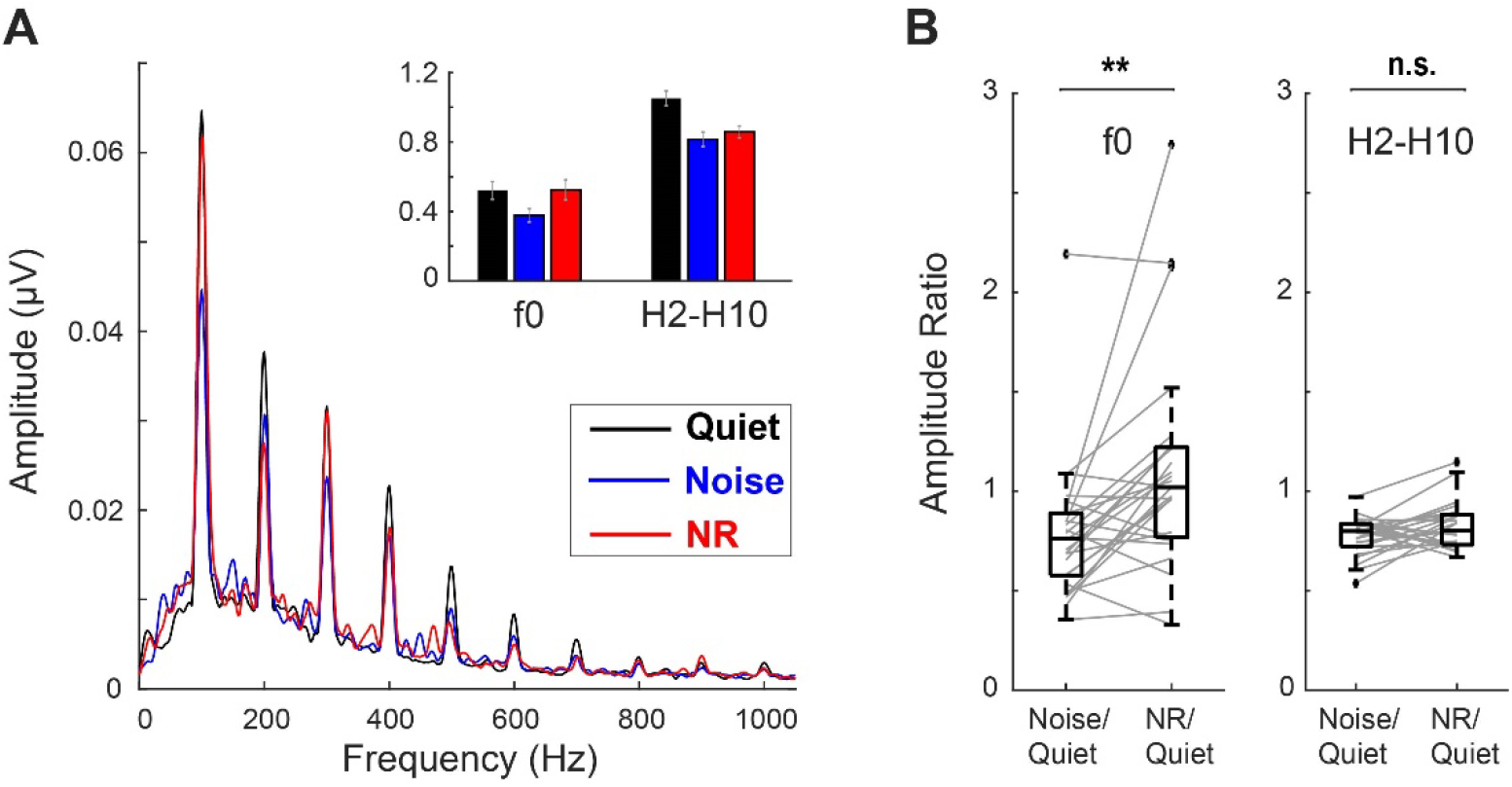
**A.** Fast Fourier transform of the vowel portions of the response (60-180 ms) with the bar graphs indicating resulting spectral amplitudes in three conditions for f0 and upper harmonics (H2-H10), respectively. **B**. The noise reduction (NR)-to-quiet amplitude ratio for f0 was significantly greater than the noise-to-quiet ratio (*t*(25) = -3.57, *p* = 0.0015), indicating that NR processing may enhance encoding of f0 in noise. ^**^: significant at *p* < 0.01, n.s.: not significant.

### 2.3 Relationship between Brainstem Measures and Behavioral Outcomes

At the individualized SNR level (−6 to -11 dB, median: -8 dB) measured through an adaptive test, behavioral accuracy was obtained in the noise condition (mean = 58.58%, SD = 7.67%). The SNR levels provided to the individual listeners were deemed adequate for targeting the midpoint (62.5%) of the psychometric function relating to SNR and accuracy, with the chance level at 25%. The mean accuracy in the NR condition was 58.31% (SD = 5.64%), which indicated that NR processing did not improve speech-in-noise performance (paired *t*-test: *t*(25) = 0.24, *p* = 0.81), consistent with the literature [e.g., Alcántara et al. (2003); Bentler et al. (2008); Ricketts and Hornsby (2005)].

Pearson correlation analyses were used to investigate whether the effects of noise and NR processing on brainstem responses predicted behavioral accuracy. A change in *r*-value from correlations between responses to the consonant (i.e., NR-to-quiet minus noise-to-quiet correlations) was not related to behavioral accuracy in the NR condition (*r*= 0.040, *p* = 0.85) (**Figure 3A** left panel) and in the noise condition (*r* = 0.17, *p* = 0.41). A change in spectral amplitude ratios for f0 (i.e., NR-to-quiet minus noise-to-quiet ratio) significantly correlated with accuracy in the NR condition (*r* = 0.48, *p* = 0.013) (**Figure 3A** right panel), whereas upper harmonics (H2-H10) did not (*r* = 0.21, *p* = 0.31). Neither the encoding of f0 (*r* = 0.36, *p* = 0.069) nor the upper harmonics (*r* = 0.073, *p* = 0.72) correlated with accuracy in the noise condition. A post hoc two-sample *t*-test showed that NR benefits in f0 encoding (i.e., NR-to-quiet minus noise-to-quiet ratio) were more salient in the better performance group (i.e., a group with accuracy in the NR condition greater than the mean: 58.31%) than the other (*t*(19.77) = 3.23, *p* = 0.0042) (**Figure 3B**). These results suggest that better encoding in f0 with NR captured by brainstem measures is associated with better behavioral outcomes with NR.

**Figure 3.**
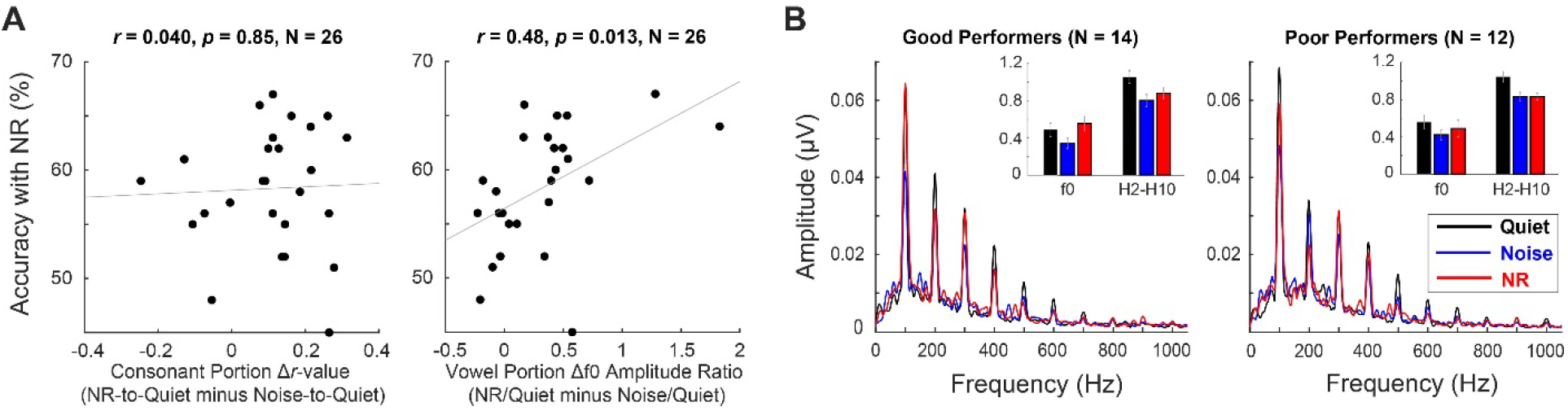
**A.** A comparison between brainstem measures and behavioral outcomes shows that the f0 amplitude ratio predicts behavioral accuracy with noise reduction (NR). **B**. Illustration of differences in spectral encoding revealed in the fast Fourier transform results between two performance groups divided based on accuracy with NR.

## 3. Discussion

The temporal and spectral metrics derived in the current study from the speech ABR were effective in capturing NR effects on subcortical speech encoding, indicating their sensitivity and feasibility. The significant correlation between the f0 amplitude ratios and behavioral accuracy with NR suggests that the individual subcortical encoding of NR-processed speech sounds influences individual behavioral outcomes related to NR processing. Our findings align well with evidence emphasizing the role of subcortical sources in accounting for individual variability in speech-in-noise perception (Bidelman & Momtaz, 2021; Gorina-Careta et al., 2021; White-Schwoch et al., 2022) and illustrating potential applications of speech-evoked ABR or envelope following response (EFR) as an objective aided outcome measure (Anderson & Kraus, 2013; Easwar et al., 2023; Easwar et al., 2015; Jenkins et al., 2018; Karawani et al., 2018). Indeed, the speech-evoked has been tested in clinical applications in recent studies with adult hearing aid users, although they did not report NR effects on brainstem responses and found no correlation between speech-evoked ABR characteristics (e.g., f0 amplitude) and behavioral outcomes (BinKhamis, Elia Forte, et al., 2019; Perugia et al., 2021; Seol et al., 2020). The current study suggests that our ABR paradigm has the potential to be an objective measure of individual noise tolerance and sensitivity to NR-processed speech and a reliable predictor of behavioral outcomes with NR.

Our findings show that individual variations in peripheral and/or subcortical physiology contribute to individual differences in preferences related to NR processing. Indeed, individual variations in the peripheral encoding of complex sounds can arise from a range of overt and hidden forms of sensorineural hearing loss (Hauser et al., 2024). For instance, in people with mild-to-moderate hearing loss who are typical candidates for hearing aid use, individual variability and suprathreshold hearing deficits may stem from reduced cochlear sensitivity and reduced frequency selectivity (Henry & Heinz, 2012; Horst, 1987; Liberman & Dodds, 1984; Moore, 2007) as well as cochlear deafferentation (Bharadwaj et al., 2014; Kujawa & Liberman, 2009), potentially leading to significant differences in subcortical encoding fidelity among individuals. Further, emerging evidence indicates that peripheral hearing damage also induces distorted cochlear tonotopy, where hypersensitive cochlear tuning curve tails allow low-frequency sounds to commandeer the temporal response of the basal half of the cochlea (Bharadwaj et al., 2024; Henry et al., 2016; Henry et al., 2019; Parida & Heinz, 2022). Lastly, although the scalp-recorded brainstem response to speech stimuli is generally assumed to be unaffected by cortical generators [see a review from Chandrasekaran and Kraus (2010)], given that NR effects in the current study were observed at 100 Hz (f0), some contribution from cortical and non-sensory cognitive variables is also possible (Forte et al., 2017; Hoormann et al., 2004; Lehmann & Schönwiesner, 2014). Future work should explore how variations in subcortical encoding interact with individual differences in cognitive processes, such as auditory selective attention (Kim, Emory, et al., 2021; Shim et al., 2023), in ultimately determining behavioral outcomes.

## 4. Methods

### 4.1 Participants

Twenty-six adults (3 male, 12%) with normal hearing participated in the experiments. All participants were native speakers of American English, with air-conduction thresholds no greater than 20 dB HL at any frequencies, tested in octaves from 250 up to 8000 Hz, and their ages ranged from 19 to 41 years (mean = 24.54 years, SD = 5.01 years). All study procedures were conducted at Purdue University and were reviewed and approved by the Purdue University Institutional Review Board. All work was carried out following the Code of Ethics of the World Medical Association (Declaration of Helsinki), and written informed consent was obtained for everyone.

### 4.2 Stimuli and Procedures

#### 4.2.1 Speech-evoked ABR Test

The current study followed well-established procedures of Parbery-Clark et al. (2009) unless otherwise noted. We used the speech syllable /da/ constructed using the KlattGrid speech synthesizer (Klatt, 1980). The speech syllable was 170 ms long with 5 ms onset and offset ramps and had an average fundamental frequency (f0) of 101 Hz (100 to 106 Hz). During the first 50 ms formant transition period, the first, second, and third formants changed from 482 to 765 Hz, 2046 to 1131 Hz, and 2695 to 2483 Hz, respectively, but stabilized for the following 120 ms steady-state (vowel) portion of the stimulus. The fourth, fifth, and sixth formants remained constant at 3626, 4060, and 5249 Hz, respectively. The acoustical characteristics described above were estimated by Praat (Boersma, 2001). Speech-shaped noise generated using a 512-order finite impulse response filter was added to the speech syllable for the noise and NR conditions.

The speech syllable was presented at 90 dB SPL monoaurally to the better ear, chosen based on pure-tone air conduction threshold averaged across 0.5, 1, 2, and 4 kHz, through insert ER-2 earphones (Etymotic Research, Elk Grove, IL). In the noise and the NR conditions, the speech syllable was presented at a +3 dB SNR over speech-shaped noise that started 40 ms before the syllable onset and continued until 40 ms after the syllable offset. Thus, the overall 250 ms long stimulus was presented in alternating polarities with an interstimulus interval of 40 ms. Speech-evoked ABR was recorded at a 16 kHz sampling rate using the BioSemi ActiveTwo 32-channel system (BioSemi B.V., Amsterdam, the Netherlands) in the international 10-20 layouts over 3000 sweeps for each of three experimental conditions (i.e., quiet, noise, and NR conditions). The magnitude of offset voltages was adjusted to be less than ±30 mV at each electrode before the beginning of data collection. Participants watched muted, captioned videos during the tests conducted in a single-walled, sound-treated booth (IAC Acoustics, Naperville, IL) through a computer monitor placed at zero-degree azimuth a half-meter distance from their eye level. The recordings spanned around 45 minutes and were controlled by custom scripts implemented in MATLAB (R2016b, the MathWorks, Natick, MA).

All EEG recordings were re-referenced to two earlobe electrodes and bandpass filtered from 70 to 2000 Hz (12 dB/octave with zero-phase shift). Trials with activity greater than ±35 µV were considered artifacts and removed from the analysis. Epochs were generated from -10 to 190 ms relative to the onset of the speech syllable /da/ and baseline-corrected relative to mean activity in the pre-syllable period. In the current study, only single-channel analyses were conducted using the Cz electrode at the vertex.

#### 4.2.2 Speech-in-noise Test

The Iowa Test of Consonant Perception was administered to assess the perception of consonant-vowel-consonant monosyllabic English words embedded in speech-shaped noise (Geller et al., 2020). Participants were tested in the same environment described above (i.e., participants tested in a sound booth, stimuli presented monoaurally through an insert ER-2 earphones, and tasks controlled in MATLAB).

Each trial started with the screen indicating a trial number in silence for 0.5 seconds, then switched to the screen with a fixation cross (‘+’) in the center to fix eye gaze during stimuli presentation to minimize eye-movement artifacts. Half a second after the cross symbol occurred on the screen, background noise started and continued for 1.5 seconds. The target word was presented 0.5 seconds after the noise onset. The composite stimulus was presented at 80 dB SPL. After the stimuli presentation, the participants were required to select one answer out of four choices shown on the screen using a keypad. For instance, for the target word *sat*, four answer options were provided: *sat, pat, fat*, and *that*. No feedback was provided during the experiment.

The current study targeted the threshold level for closed-set tests (i.e., 62.5% accuracy, halfway between the chance level of 25% and 100%). Kim et al. (2022) used an adaptive test with the two-down, one-up staircase procedure to find the SNR level targeting 70% accuracy (i.e., speech reception threshold, SRT-70) in the first 50 trials and reported that in listeners with normal hearing, the SNR level 3-dB lower than the SRT-70 led to mean accuracy of 62.4%. The present study followed the procedures from Kim et al. (2022); for the first 50 trials, the adaptive test was used to find the SRT-70 for individual listeners, and then the SNR level 3-dB lower than the SRT-70 was given to them in the following two experimental conditions: the noise and NR conditions. One hundred ten words were presented in each condition using word sets balanced across speaker gender and initial phonemes and randomly assigned to each condition.

### 4.3 NR Algorithm

In the NR conditions for both the speech-evoked ABR test and speech-in-noise test, the current study used the Ephraim-Malah NR algorithm, a modified spectral-subtraction NR that applies different gain across frequency channels in each short-time frame based on SNR estimation using a minimum mean-square error estimator (Ephraim & Malah, 1984). The Ephraim-Malah NR algorithm has relatively low computational complexity and thus has been implemented in modern digital hearing aids [e.g., Sarampalis et al. (2009); Stelmachowicz et al. (2010)].

**Figure 4** upper panel illustrates spectra of speech and noise stimuli from the speech-evoked ABR tests extracted using the phase-inversion technique (Hagerman & Olofsson, 2004): inverting the noise phase before processing two noisy signals through NR. The NR algorithm in the present study improved the SNR by approximately 2.5 dB based on the long-term RMS level of those extracted speech and noise stimuli. **Figure 4** lower panel shows magnitude-squared coherence up to 8 kHz for two speech stimuli (i.e., unprocessed vs. NR-processed speech stimuli extracted from the phase-inversion technique) where coherence value zero indicates that input and output power spectra are not identical at all, whereas value one means they are entirely identical (Kay, 1988). The coherence value averaged across 8 kHz was at around 0.3 between speech stimuli from the speech-evoked ABR tests. For monosyllabic words used for speech-in-noise tests in the present study, approximately 3.5-dB benefit in SNR and average coherence at around 0.6 were reported by Kim et al. (2022).

**Figure 4.**
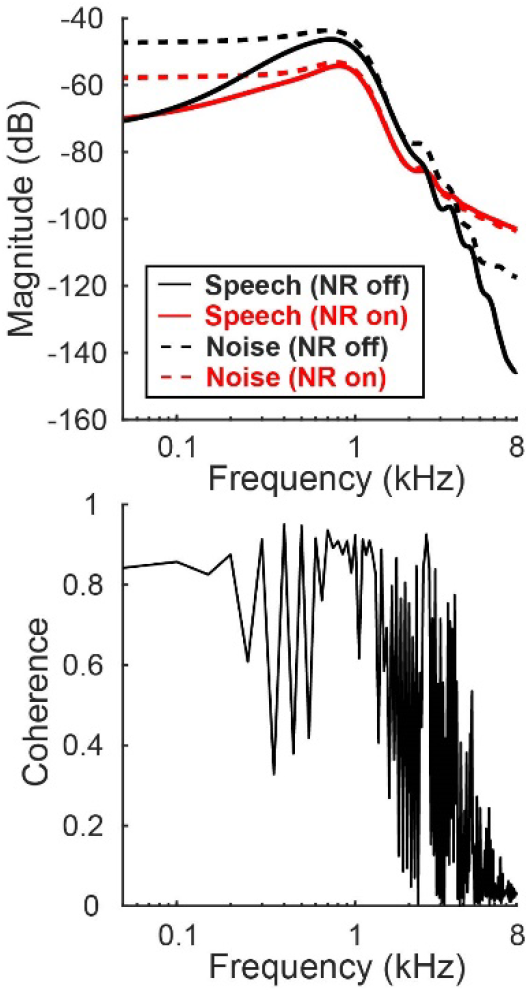
The upper panel shows spectra of speech and noise stimuli for the speech-evoked auditory brainstem response (ABR) tests. Noise reduction (NR)-processed stimuli were extracted using the phase-inversion technique. The lower panel illustrates the magnitude-squared coherence across frequencies between unprocessed and NR-processed speech stimuli for the ABR tests.

### 4.3 Statistical Analysis

#### Response-to-response Correlations

Cross-correlation analyses were conducted to examine the effects of noise and NR on the consonant portion of brainstem responses (5-60 ms). Specifically, brainstem responses in both NR and noise conditions were compared to the responses in quiet, respectively. Given that adding noise or NR may delay the responses relative to the responses in quiet, the response waveforms in noise and NR conditions were shifted in time by up to 2 ms until reaching the maximum correlation coefficient (a Pearson’s r value) (Parbery-Clark et al., 2009). A pair of correlation coefficients were compared: noise-to-quiet vs. NR-to-quiet.

#### Spectral Encoding

Fast Fourier transform analysis was conducted on the vowel portions of the response (60-180 ms) in the same manner as described by Parbery-Clark et al. (2009) to assess the effect of noise on spectral encoding to the vowel portions of the speech syllable and to examine if NR processing enhances spectral encoding in noise. The strength of harmonic representation (100-1000 Hz) with the first harmonic corresponding to the stimulus f0 (100 Hz) was quantified by summing spectral amplitudes across frequency bins that are 20-Hz wide and centered on each of the ten harmonics. Aside from calculating the f0 spectral amplitude, spectral amplitudes for the 2nd to 10th harmonics were combined to represent the strength of overall spectral encoding to the subsequent harmonics. NR-to-quiet and noise-to-quiet amplitude ratios were calculated and compared for f0 and upper harmonics, respectively.

#### Relationship between Brainstem Measures and Behavioral Outcomes

The relationship between the brainstem measures described above, correlations between responses to the consonant and spectral amplitude ratios for the vowel portion of the response, and behavioral outcomes were assessed using Pearson correlation analysis. Specifically, the correlation analyses investigated if a change in *r*-value from response-to-response metrics and a change in spectral amplitude ratios correlate with behavioral accuracy, respectively. A post hoc two-sample *t*-test was conducted to compare NR benefits (or lack thereof) in f0 encoding between two participant groups divided based on accuracy with NR: one group with accuracy greater than the mean vs. the other group with accuracy less than the mean.

## Data Availability

All data is openly accessible and publicly available at https://dx.doi.org/10.17632/5jgbmk6p42.1.

## Code Availability

The code to reproduce the figures in this work is publicly available at https://dx.doi.org/10.17632/5jgbmk6p42.1.

## Acknowledgements

This work was supported by the Hearing Health Foundation (Emerging Research Grant 2022 & 2023, PI: Kim), Royal National Institute for Deaf People (Flexi Grant F110, PI: Kim), and NIH/NIDCD (R01DC015989, PI: Bharadwaj).

## Author Contributions

S.K. and H.M.B. designed the experiments and analyzed the data. S.K. and M.S. ran the experiments. All authors wrote the manuscript.

## Competing Interests

The authors declare no competing interests.

## Notes

### Competing Interest Statement

The authors have declared no competing interest.

